# Improved prediction of protein-protein interactions using AlphaFold2

**DOI:** 10.1101/2021.09.15.460468

**Authors:** P. Bryant, G. Pozzati, A. Elofsson

**Affiliations:** Science for Life Laboratory, 172 21 Solna, Sweden; Department of Biochemistry and Biophysics, Stockholm University, 106 91 Stockholm, Sweden

**Author notes:** These authors contributed equally: P. Bryant, G. Pozzati.

## Abstract

Predicting the structure of interacting protein chains is a fundamental step towards understanding protein function. Unfortunately, no computational method can produce accurate structures of protein complexes. AlphaFold2, has shown unprecedented levels of accuracy in modelling single chain protein structures. Here, we apply AlphaFold2 for the prediction of heterodimeric protein complexes. We find that the AlphaFold2 protocol together with optimized multiple sequence alignments, generate models with acceptable quality (DockQ≥0.23) for 63% of the dimers. From the predicted interfaces we create a simple function to predict the DockQ score which distinguishes acceptable from incorrect models as well as interacting from non-interacting proteins with state-of-art accuracy. We find that, using the predicted DockQ scores, we can identify 51% of all interacting pairs at 1% FPR. The protocol can be found at: https://gitlab.com/ElofssonLab/FoldDock.

## Introduction

Protein-protein interactions are central mediators in biological processes. Most interactions are governed by the three-dimensional arrangement and the dynamics of the interacting proteins^1^. Such interactions vary from being permanent to transient^2,3^. Some protein-protein interactions are specific for a pair of proteins, while some proteins are promiscuous and interact with many partners. This complexity of interactions is a challenge both for experimental and computational methods.

Often, studies of protein-protein interactions can be divided into two categories, the identification of what proteins interact and the identification of how they interact. Although these problems are distinguished, some methods have been applied to both problems ^4,5^. Protein docking methodologies refer to how proteins interact and can be divided into two categories considering proteins as rigid bodies; those based on an exhaustive search of the docking space^6^ and those based on alignments (both sequence and structure) to structural templates^7^. Exhaustive approaches rely on generating all possible configurations between protein structures or models of the monomers^8,9^ and selecting the correct docking through a scoring function, while template-based docking only needs suitable templates to identify a few likely candidates. However, flexibility has often to be considered in protein docking to account for interaction-induced structural rearrangements^10,11^. Therefore, flexibility limits the accuracy achievable by rigid-body docking^12^, and flexible docking is traditionally too slow for large scale applications. A possible compromise is represented by semi-flexible docking approaches^13^ that are more computationally feasible and can consider flexibility to some degree during docking.

Regardless of different strategies, docking remains a challenging problem. In the CASP13-CAPRI experiments, human group predictors achieved up to 50% success rate for topranked docking solutions^14^. Alternatively, a recent benchmark study^8^ reports success rates of different web-servers reaching up to 16% on the well known Benchmark 5 dataset^15^.

Recently, in the CASP14 experiment, AlphaFold2 (AF2) reached an unprecedented performance level in structure prediction of single-chain proteins^16^. Thanks to an advanced deep learning model that efficiently utilises evolutionary and structural information, this method consistently outperformed all competitors, reaching an average GDT_TS score of 90^16^. Recently, RoseTTAFold was developed, trying to implement similar principles^17^. Since then, other end-to-end structure predictors have emerged using different principles such as fast multiple sequence alignment (MSA) processing in DMPFold2^18^ and language model representations^19^.

As an alternative to other docking methods, it is possible to utilise co-evolution to predict the interaction between two protein chains. Initially, direct coupling analysis was used to predict the interaction of bacterial two-component signalling proteins ^20,21^. Later, these methods were improved using machine learning^22^.

In a Fold and Dock approach, two proteins are folded and docked simultaneously. We recently developed a Fold and Dock pipeline using another distance prediction method focused on protein folding (trRosetta^23^). In this pipeline, the interaction between two chains from a heterodimeric protein complex and their structures were predicted using distance and angle constraints from trRosetta^24,25^. This study demonstrated that a pipeline focused on intra-chain structural feature extraction can be successfully extended to derive inter-chain features as well. Still, only 7% of the tested proteins were successfully folded and docked.

In that study, we found that generating the optimal MSA is crucial for obtaining accurate Fold and Dock solutions, but this is not always trivial due to the necessity to identify the exact set of interacting protein pairs^26^. Given the existence of multiple paralogs for most eukaryotic proteins, this is difficult. We also found that this process requires an optimal MSA depth to optimise inter-chain information extraction. Too deep MSAs might contain false positives (i.e. protein pairs that interact differently), resulting in noise masking the sought after co-evolutionary signal, while too shallow alignments do not provide sufficient co-evolutionary signals.

In this work we systematically apply the AF2 pipeline on two different datasets to Fold and Dock protein-protein pairs simultaneously. We explore the docking success using the AF2 pipeline in combination with different input MSAs, in order to study the relationship between the output model quality and these inputs. We also find that, by scoring multiple models of the same protein-protein interaction with a predicted DockQ score (pDockQ), we can distinguish with high confidence acceptable (DockQ≥0.23) from incorrect models. The modelling success is higher for bacterial protein pairs, pairs with large interaction areas consisting of helices or sheets, and many homologous sequences. We also test the possibility to distinguish interacting from non-interacting proteins and find that, using pDockQ, we can separate truly interacting from non-interacting proteins with consistent accuracy. We find that the results in terms of successful docking using AF2 are superior to other docking methods. AF2 clearly outperforms a recent state-of-the-art method^27^ and our protocol performs quite close to (63% vs 72%) the recently developed AF-multimer^28^, which was developed using the same data as the test set here, making a direct comparison difficult.

## Results and Discussion

### Identifying the best AlphaFold2 model

The success rate (SR), i.e. the percentage of acceptable models (DockQ>0.23), is used to measure AF2 performance over the development set using the different multiple sequence alignment (MSAs). The best performance is 33.3% for the AF2 MSAs and 39.4% for the AF2+paired MSAs (Table 1). It is thereby evident that combining both paired and AF2 MSAs is superior to using them separately. The average performance of the AF2 and the paired MSAs is similar, but for individual protein pairs, frequently one of the two MSAs is superior to the other, as seen from that the Pearson correlation coefficient for the DockQ scores between AF2 vs paired MSAs is 0.54 (Table S2). Therefore, combining AF2 and paired MSAs improves the results.

**Table 1.** Success rate of different modelling setups.

| Table 1. Success rate of different modelling setups |  |  |  |  |
| --- | --- | --- | --- | --- |
| Neural network configuration |  |  |  |  |
| NN model | model_1 | model_1 | model_1_ptm | model_1_ptm |
| Recycles | 10 | 3 | 10 | 3 |
| Ensembles | 1 | 8 | 1 | 8 |
| Setup short name | m1-10-1 | m1-3-8 | mp-10-1 | mp-3-8 |
| Paired MSA | **28.7** | 28.2 | **28.7** | 27.8 |
| AF2 MSA | 31.5 | **33.3** | 26.4 | 23.6 |
| AF2+Paired MSA | **39.4** | 38.4 | 32.4 | 31.0 |

Results of AF2 run on the development set using different MSAs and neural network configurations. Row labels in bold indicate AF2 setup features. Every column in the table refers to an overall setup and every corresponding value refers
to a run of the described setup with a different input MSA. Values represent percentage of acceptable models (DockQ ≥ 0.23) overall the development set. The highest success rates for each MSA type are highlighted in bold.

Next, we compared the default AF2 model (model_1) with the fine-tuned versions of (model_1_ptm). Surprisingly, the original AF2 model_1 outperforms AF2 model_1_ptm in most cases (Table 1). Further, the difference between 10 recycles-one ensemble and three recycles-eight ensembles is minor across all MSAs and AF2 models. Therefore, the input information and the AF2 model appears to impact the outcome the most.

### Test set performance

The best model and configuration for AF2 (m1-10-1) was used for further studies on the test set. The best outcome using this modelling strategy results in an SR of 57.8% (856 out of 1481 correctly modelled complexes) for the AF2+paired MSAs compared with 45.0% using the AF2 MSAs alone (Figure 1, Table 2). The results using the block diagonalization+paired MSAs are almost identical (SR=58.4%, median=0.363). Further, running five initialisations with random seeds and ranking the models using the predicted DockQ score (pDockQ, Figure 2C), increases the SR to 61.7% and 62.7% for the AF2+paired and block diagonalization+paired MSAs, respectively (model variation and ranking, Figure 2). Using the combination of AF2 and paired MSAs increases performance, suggesting that AF2 gains both from larger and paired MSAs, although it often can manage with less information.

**Figure 1.**
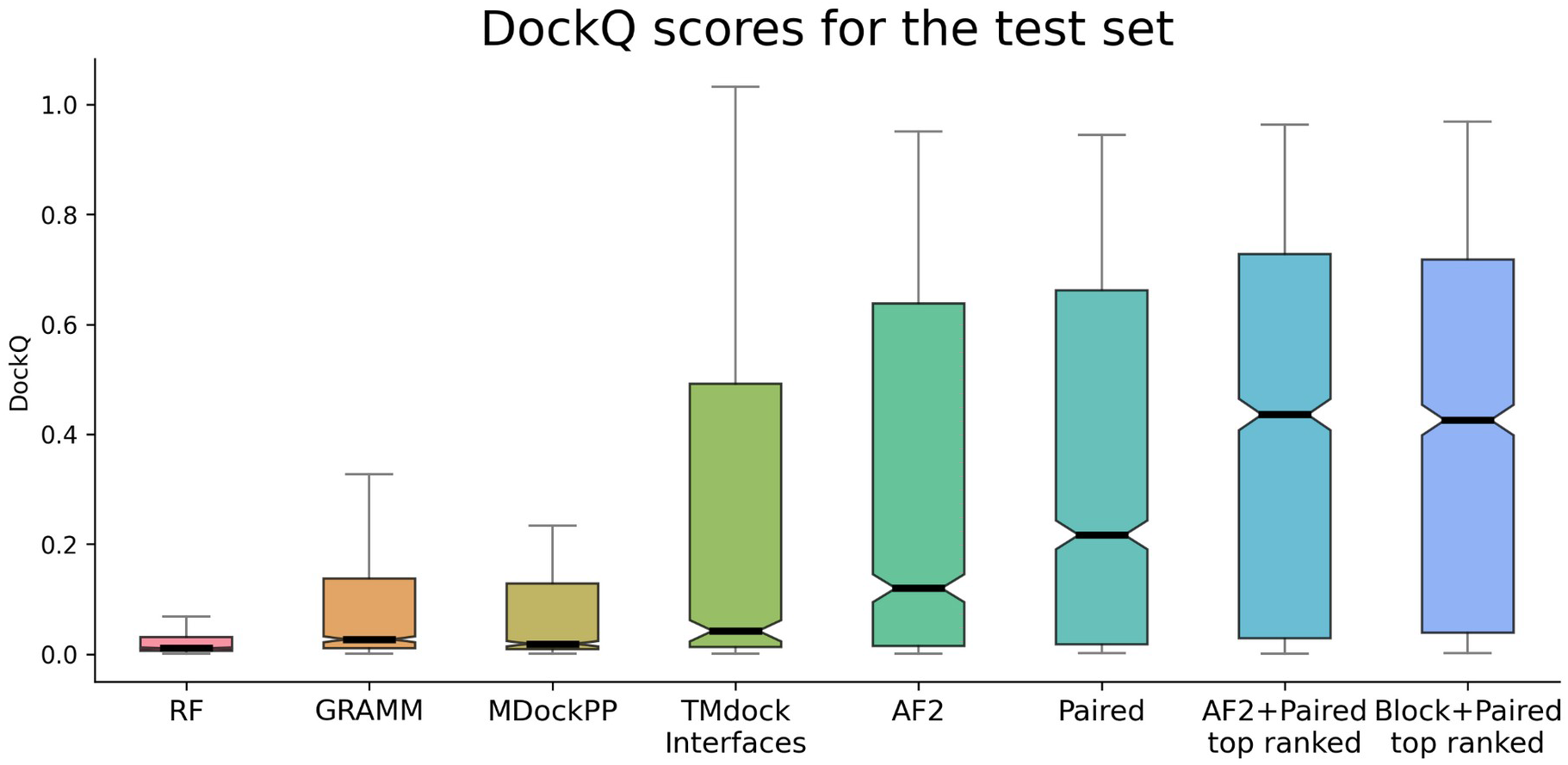
DockQ scores for the test set. Distribution of DockQ scores as boxplots for different modelling strategies on the test set. Boxes encompass data quartiles, horizontal lines mark the medians and upper and lower whiskers indicate respectively maximum and minimum values for each distribution. All AF2 models have been run with the same neural network configuration (m1-10-1). Outlier points are not displayed here. AF2, refers to running AF2 using the default AF2 MSAs, “Paired” refers to using MSAs paired using information about species and “Block” refers to using block diagonalization MSAs.

**Figure 2.**
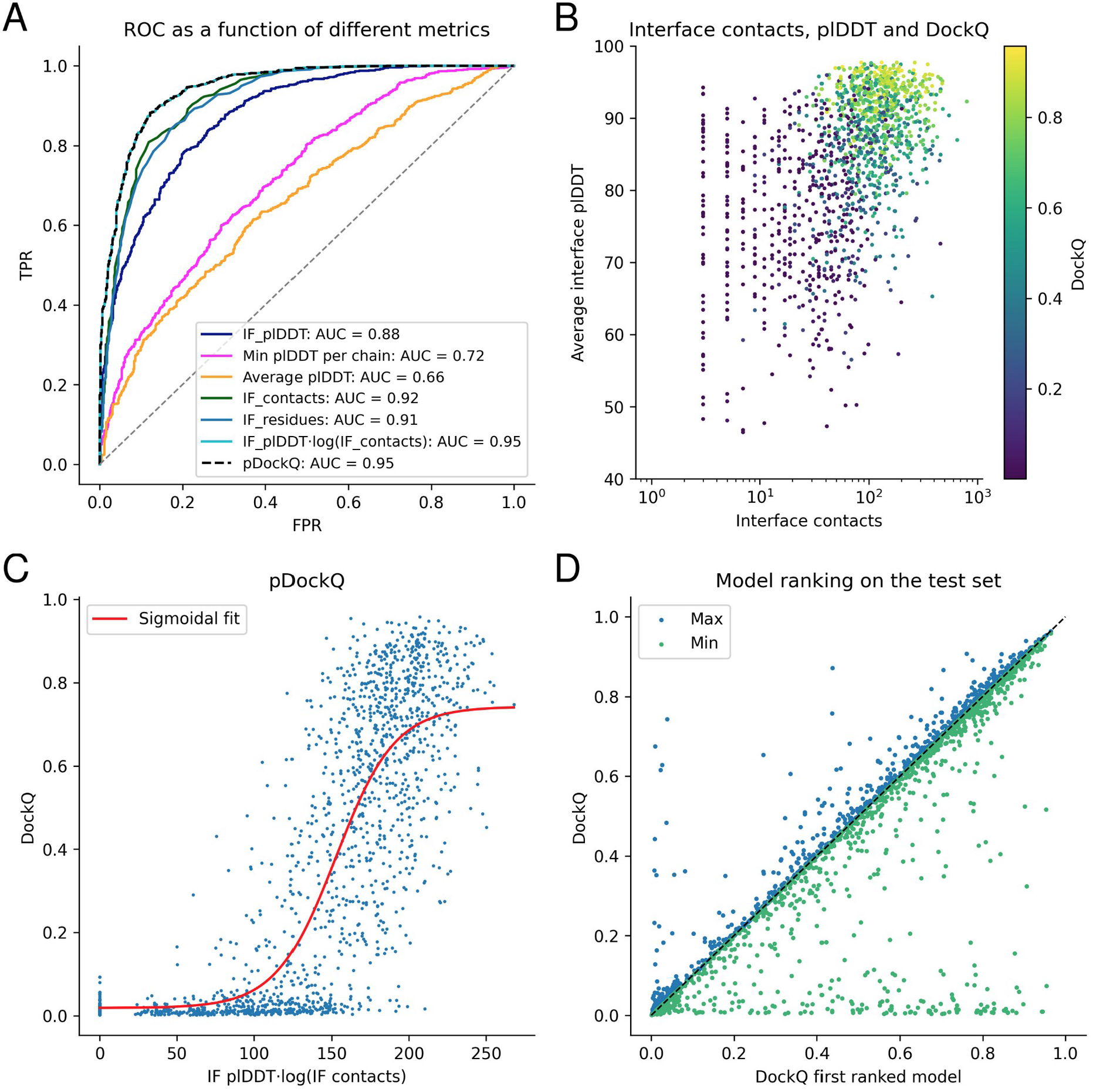
Model quality metrics and multiple model ranking. **A)** ROC curve as a function of different metrics for the test dataset (first run). Cβs within 8 Å from each other from different chains are used to define the interface. IF_plDDT is the average plDDT of interface residues, min plDDT per chain is the minimum average plDDT of both chains, average plDDT is the average of the entire complex and IF_contacts and IF_residues are the number of interface residues and contacts respectively. pDockQ is a sigmoidal fit to the combined metric IF_plDDT·log(IF_contacts) fitted to predict DockQ as the target score, see C. **B)** Average interface plDDT vs log interface contacts colored by DockQ score on the test set. Increasing both the number of interface contacts and average interface plDDT results in higher DockQ scores. **C)** Using the combined metric IF_plDDT·log(IF_contacts), we fit a sigmoidal curve towards the DockQ scores on the test set, enabling predicting the DockQ score in a continuous manner (pDockQ). The average error overall is 0.14 DockQ score. **D)** Impact of different initialisations on the modelling outcome in terms of DockQ score on the test dataset. The maximal and minimal scores are plotted against the top-ranked models using the pDockQ scores for the AF2+paired MSAs, m1-10-1.

**Table 2.** Success rate and median DockQ scores for the test set using different methods and model configurations. “Block” refers to block diagonalization MSAs.

|  |  |  |
| --- | --- | --- |
| **Method** | **Success Rate (%)** | **Median DockQ** |
| RoseTTAfold | 9.6 | 0.011 |
| GRAMM | 21.4 | 0.027 |
| MDockPP | 24.2 | 0.019 |
| TMdock | 33.6 | 0.040 |
| TMdock interfaces | 35.1 | 0.042 |
| AlphaFold2 | 45.0 | 0.120 |
| Paired | 49.6 | 0.217 |
| AlphaFold2+Paired | 57.8 | 0.382 |
| Block+Paired | 58.4 | 0.363 |
| AlphaFold2+Paired top ranked | 61.7 | 0.436 |
| Block+Paired top ranked | 62.7 | 0.426 |
| AF-multimer | 72.2% | 0.560 |

What is most striking is that AF2 outperforms all other tested docking methods by a large margin (Figure 1, Table 2). RF is better than AF2 only for 14 pairs in the test set, while GRAMM and template-based docking (TMdock interface) outperform AF2 for 188 and 225 pairs, respectively. The best performing method in the CASP14-CAPRI experiment^29^, MDockPP^30^, achieves a SR of only 24.2%. The reason for GRAMM’s, TMdock’s and MDockPP’s good performance is likely due to the use of the bound form of the proteins, resulting in very high shape complementarity and therefore having the “answer” provided in a way.

The recently developed AF-multimer^28^ has the best performance (SR=72.2%, median=0.560, Table 2). This method is developed using the same data as the test set, which makes a direct comparison difficult. Regardless, we do believe it is likely that using AF-multimer, the performance would increase over the results of our pipeline, but it is possible the difference is less than the observed 9%.

### Distinguishing acceptable from incorrect models

It is not only essential to obtain improved predictions, but also to be able to discriminate between acceptable and non-acceptable ones. We measure the separation between correct (DockQ≥0.23) and incorrect models provided by several metrics using a receiver operating characteristic (ROC) curve. Different criteria were examined over the test set, including (i) the number of unique interacting residues (Cβ atoms from different chains within 8 Å from each other) in the interface, (ii) the total number of interactions between Cβ atoms in the interface, (iii) the average plDDT for the interface, (iv) the lowest plDDT of each single chain average, and (v) the average plDDT over the whole protein heterodimer (Figure 2A). Three criteria result in very similar areas under the curve (AUC) measures. The total number of interactions between Cβs and the number of residues in the interface can separate the correct/incorrect models with an AUC of 0.92 and 0.91 respectively, while the average interface plDDT results in an AUC of 0.88. However, pLDDT results in higher TPRs at lower FPRs; therefore, we multiply the plDDT with the logarithm of the interface contacts resulting in an AUC of 0.95.

Interestingly, the average plDDT of the entire complex only results in an AUC of 0.66, suggesting that both single chains in a complex are often predicted very well, while their relative orientation may still be incorrect.

Figure 2B shows that increasing both the number of interface contacts and the average interface plDDT results in higher DockQ scores for the test set. Using the combination of plDDT with the logarithm of the interface contacts, we therefore fit a simple sigmoidal function to the DockQ scores (Figure 2C), see methods. This enables the prediction of the DockQ scores (pDockQ) in a continuous manner with an overall average error of 0.11 on the test set. The AUC using pDockQ as a separator is identical to the combination of plDDT with the logarithm of the interface contacts, 0.95 (Figure 2A).

### Model variation and ranking for the test set

Five models are generated using the best strategy (m1-10-1 with AF2+paired MSAs) with different initialisation (random seeds). The average SR (57.2% ± 0.0%) is similar for all five runs. However, the average deviation for individual models is DockQ=0.08 when comparing the best and worst models for a target (Figure 2D), i.e. there is some randomness to the success for an individual pair. If the maximal DockQ score across all models is used, the SR would be 62.9%. Although this is unachievable, ranking the models using the pDockQ score results in an SR of 61.7%. The AUC using the same metric for the ranked test set is 0.93, which means that 31% of all models are acceptable at an error rate of 1% and 54% at an error rate of 10% (Table S3).

### Bacterial complexes are predicted more accurately

In the test set, about 60% of the complexes can be modelled correctly. We try to identify what distinguishes the successful and unsuccessful cases by analysing different subsets of the test set. First, we divide the proteins by taxa, next by interface characteristics and finally by examining the alignments.

The Success Rates (SRs) for each kingdom is; Eukarya 61%, Bacteria 73.7%, Archaea 84.5%, and Virus 60% (Figure S1B). Further, the SRs for *S.cerevisiae* is better than for *Homo Sapiens* (66% vs 58%, Figure 3D). The higher performance in prokaryotes is consistent with previous observations regarding the availability of evolutionary information in prokaryotes compared to Eukarya^27^ (Figure S1). The higher performance in *S.cerevisiae* compared to *Homo Sapiens* suggests a similar relationship between higher and lower order organisms within the same kingdom.

**Figure 3.**
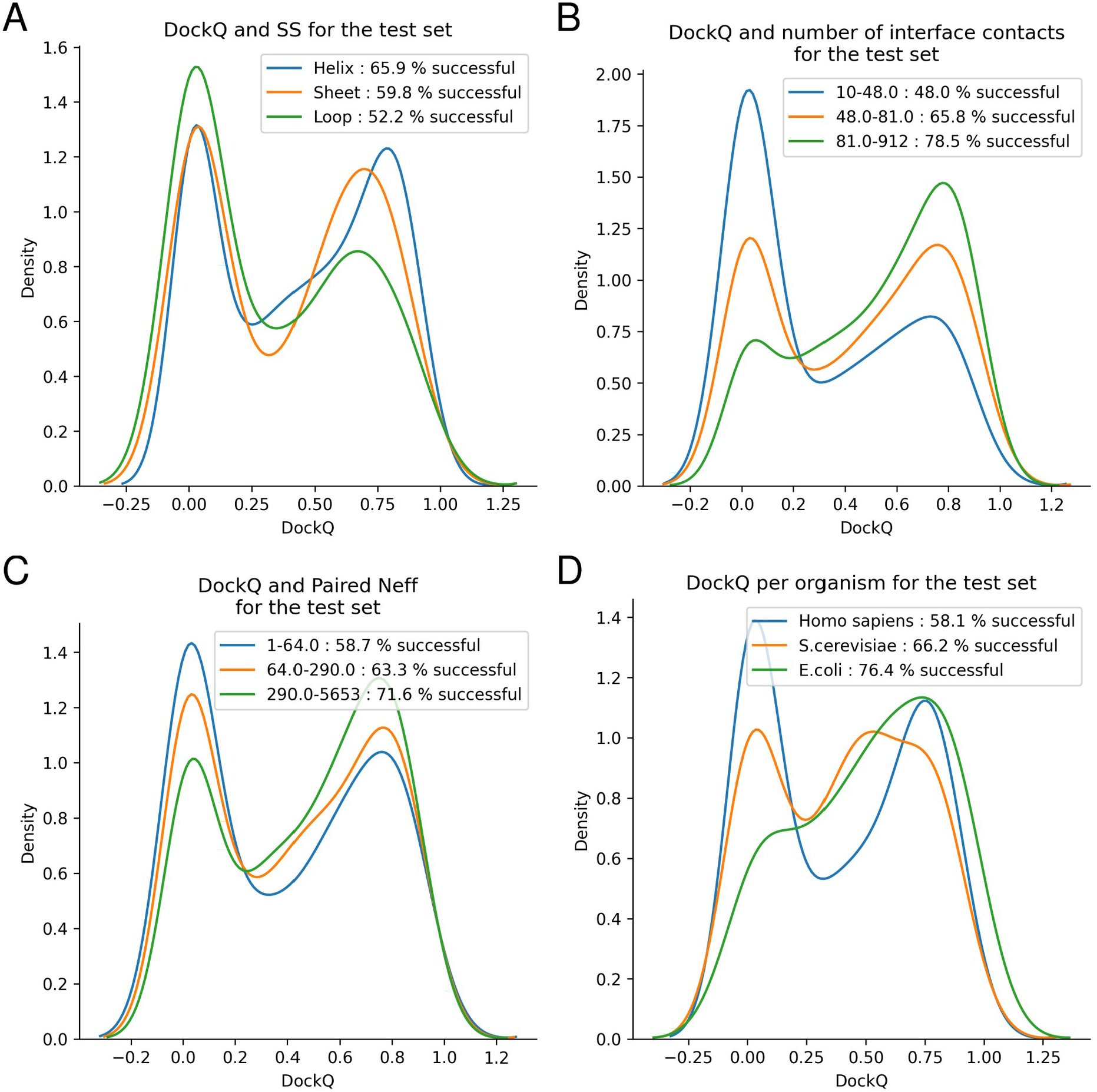
DockQ distributions for test dataset tertiles. **A)** Distribution of DockQ scores for three sets of interfaces with the majority of Helix, Sheet and Coil secondary structures. **B)** Distribution of DockQ scores for tertiles derived from the distribution of contact counts in docking model interfaces. **C)** Distribution of DockQ scores for tertiles derived from the distribution of Paired MSAs Neff scores. **D)** Distribution of DockQ scores for the top three organisms *Homo Sapiens, S. cerevisiae* and *E. coli.*

Next, we examine the interfaces. Different secondary structural content of the native interfaces is investigated (Figure 3A). The highest SR is obtained mainly for helix interfaces (62%), followed by interfaces containing mainly sheets (59%). The loop interface SR of 53% is substantially lower than the others, suggesting that interfaces with more flexible structures are harder to predict. We divide the dataset by interface size, and find that pairs with larger interfaces are easier to predict, as the SR increases from 47 to 74% between the smallest and biggest tertiles (Figure 3B).

We continue to examine features of the MSAs. First, the impact of the number of non-redundant sequences (Neff) in both paired and AF2 MSAs was analysed. It is clear that the fraction of correctly modelled sequences increases with larger Neff scores (Figure 3C). Also, paired MSA Neff (Figure 3C) has a stronger influence on the outcome than the Neff of the AF2 MSAs (Figure S2A). Secondly, the MSA interface signal in the paired MSAs, measured by the fraction of correct interface contacts using direct coupling analysis (DCA), was analysed. MSAs with stronger interface signals show higher success rates, even if the paired MSAs are used in combination with the AF2 MSAs (Figure S3). This suggests that MSA co-evolutionary signal and, thereby, correct identification of orthologous protein sequences, has a strong impact on the outcome.

### CASP14 and novel proteins without templates

Chains derived from CASP14 heteromeric targets and chains from PDB complexes with no templates are folded in pairs using the presented AF2 pipeline (default AF2+paired MSAs, ten recycles, m1-10-1 and five differently seeded runs).

For the CASP14 chains, four out of six pairs display a DockQ score larger than 0.23 (SR of 67%). No ranking is necessary in this case, given that all produced docking models for the same chain pair are very similar (the average standard deviation is 0.01 between each set of DockQ scores). An interesting unsuccessful docking is obtained modelling chains from the complex with PDB ID 6TMM (Figure S4), which are known to form a heterotetramer. In this structure, each chain A is in contact with its partner chain B at two different sites. Both docking configurations (6TMM_A-B and 6TMM_A-D) put the chain in between the two binding sites. The other unsuccessful docking (6VN1_A-H) has an interface of just 19 residue pairs.

The SR for docking the proteins without templates is 50%. Between the five different initialisations, the average difference in the DockQ score is 0.03, and there is no deviation in SR, i.e. ranking did not improve the SR. Two acceptable models are displayed in Figures 5A (7EIV_A-C) and B (7MEZ_A-B). More interesting, in one of the incorrect models (7NJ0_A-C, Figure S5), the predictions get the location of both chains correct, but their orientations wrong, resulting in DockQ scores close to 0. For 7EL1_A-E (Figure 4C), the shorter chain E is not folded correctly, and instead of folding to a defined shape, it is stretched out and inserted within chain A. It occupies the shape of the DNA in the native structure. In the two remaining incorrect models (7LF7_A-M and 7LF7_B-M), Figure 4D, the chains only interact with a short loop of the M chain, making the docking very difficult and possibly biologically meaningless.

**Figure 4.**
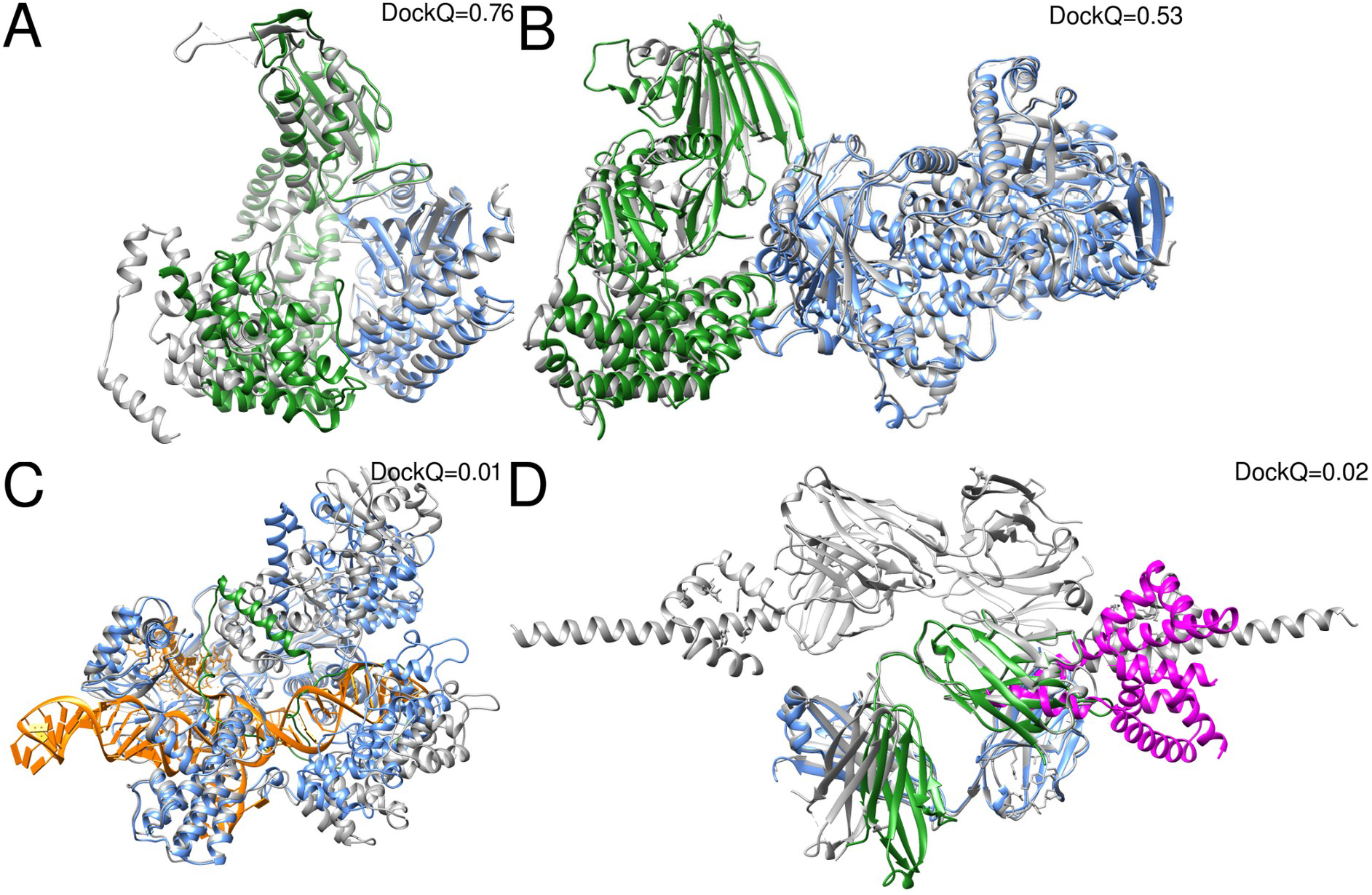
Predicted and native structures from the set of novel proteins without templates. The native structures are represented as grey ribbons **A)** Docking of 7EIV chains A (blue) and C (green) (DockQ=0.76). **B)** Docking of 7MEZ chains A (blue) and B (green) (DockQ=0.53). **C)** Prediction of structure 7EL1 chains A (blue) and E (green) (DockQ=0.01). The DNA going through chain A is coloured in orange. **D)** Docking of 7LF7 chains A (blue) and M (magenta) (DockQ=0.02) and chains B (green) and M (magenta) (DockQ=0.02).

**Figure 5.**
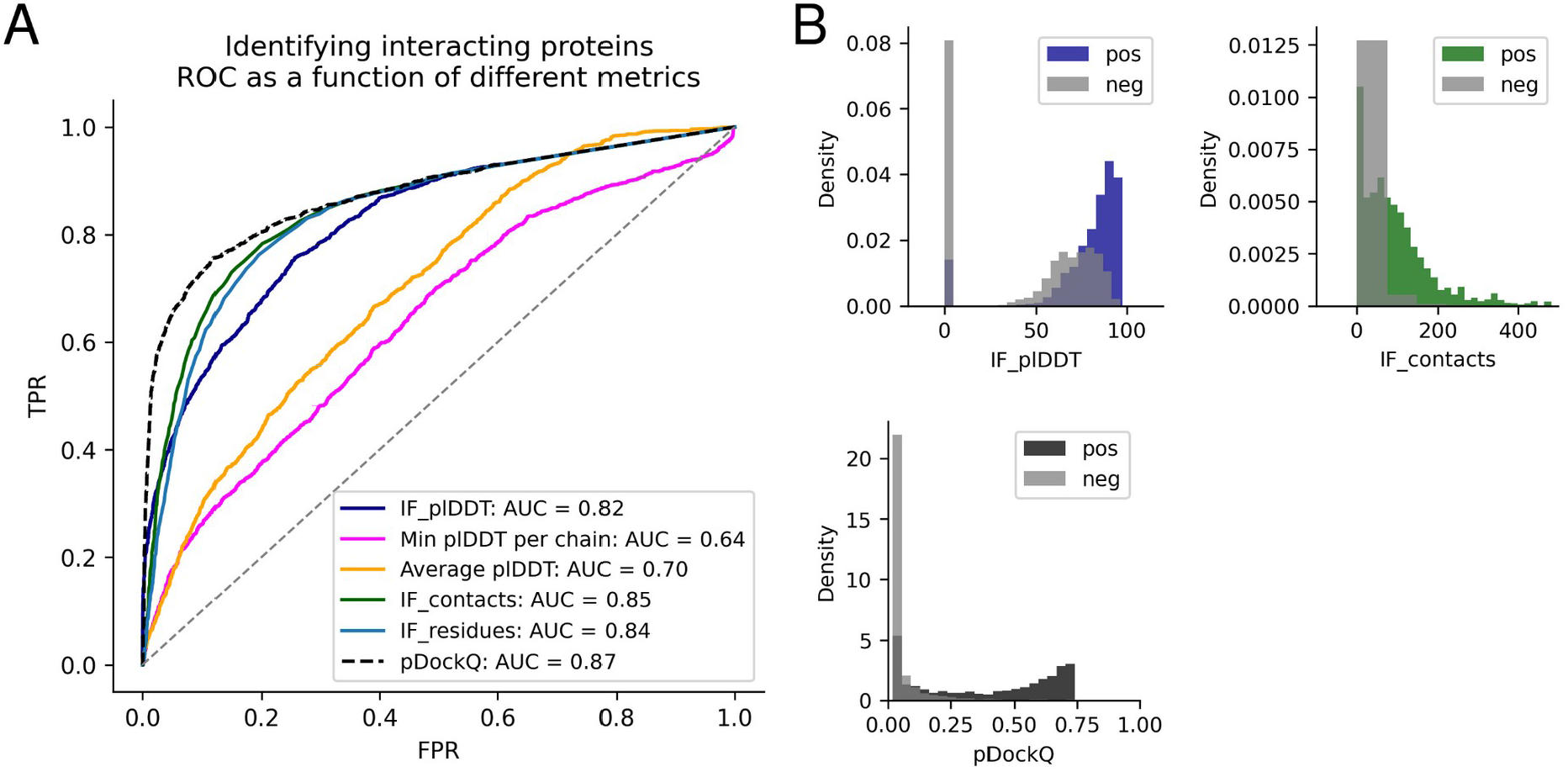
Discrimination of interacting and non-interacting proteins. **A)** The ROC curve as a function of different metrics for discriminating between interacting and non-interacting proteins. IF_plDDT is the average plDDT in the interface, min plDDT per chain is the minimum average plDDT of both chains, average plDDT is the average of the entire complex and IF_contacts and IF_residues are the number of interface residues and contacts respectively. pDockQ is a sigmoidal fit to this with DockQ as the target score, as described above. **B)** Distribution of the top discriminating features average interface plDDT (IF_plDDT), the number of interface contacts (IF_contacts), the combination of these (IF_plDDT·log(IF_contacts)) and the pDockQ for interacting (non-grey) and non-interacting proteins (grey).

### Identifying interacting proteins

Using the best separator from the model ranking, the pDockQ, it is possible to distinguish the 3989 non-interacting proteins from *E.coli* and the truly interacting proteins from the test set with an AUC of 0.87. Another recently published method obtains AUC 0.76 on this set^27^. However, these results are probably overstated since the negative set only contains bacterial proteins, while the positive set is mainly eukaryotic.

To obtain a more realistic estimate, we also include a set of non-interacting proteins from mammalian organisms combined with the non-interacting proteins from *E.coli.* On this set, we obtain an AUC of 0.82 for the average interface plDDT and slightly higher (0.84 and 0.85) for the number of interface contacts and residues, respectively (Figure 5A). pDockQ results in an ROC curve with an AUC of 0.87. Importantly, pDockQ provides a better separation at low FPRs, enabling a TPR of 51% at FPR of 1% compared to 27%, 18 and 13% for the interface plDDT, number of interface contacts and residues, respectively. At FPR 5%, the number of interface contacts and residues report TPRs of 49 and 42%, respectively, compared to 43% for the average interface plDDT and 66% for pDockQ. The distribution of the top separators can be seen in Figure 5B.

### Limitations

Here, we only consider the structures of protein complexes in their heterodimeric state, although each protein chain in these complexes may have homodimer configurations or other higher-order states. It is also possible that the complex itself exists as part of larger biological units, in potentially more complex conformations. Investigating alternative oligomeric states and larger biological assemblies is outside of the scope of this analysis and left for future work.

The study of AF2s ability to separate interacting and non-interacting proteins here contains more extensive data than recent studies^27^. However, to test this separation thoroughly, the data studied here needs to be extended to compare interactions within individual organisms. We leave this extensive analysis to further studies.

### Findings and future prospects

Here we show that AlphaFold2 (AF2) can predict the structure of many heterodimeric protein complexes, although it is trained to predict the structure of individual protein chains. Even using the default settings, it is clear that AF2 is superior to all other tested docking methods, including other Fold and Dock methods^17,24^, methods based on shape complementarity^31,30^ and template-based docking. Using optimised multiple sequence alignments with AF2, we can accurately predict the structure of heterodimeric complexes for an unprecedented success rate of 62.7% on a large test set. The success rate is higher in E.Coli (76.4%) than in *Homo Sapiens* or *S. cerevisiae* (58.1 and 66.2% respectively).

Further, by analyzing the predicted interfaces, we can predict the DockQ score (pDockQ) with an average error of 0.1, resulting in the separation of acceptable and incorrect models with an AUC of 0.95. This means that 31% of the models can be called acceptable at a specificity of 99% (or 54% at 90% specificity). Interestingly, no additional constraints are implemented in AF2 to pull two chains in contact, meaning chain interactions (and subsequently interface sizes) are exclusively determined by the amount of inter-chain signals extracted by the predictor. Assuming that all residues in an interface contribute to the interaction energy could explain why larger interfaces are more likely to be correctly predicted.

We find that the MSA generation process can be sped up substantially at no performance loss (performance increase of 1 % SR) by simply fusing MSAs from two HHblits runs on Uniclust30 instead of using the MSAs from AF2. Fast MSA generation circumvents the main computational bottleneck in the pipeline. Using pDockQ makes it possible to separate truly interacting from non-interacting proteins with an AUC of 0.87, making it possible to identify 51% of interacting proteins at an error rate of 1%. The pDockQ score discriminates between both model quality and binary interactions. Therefore, the same pipeline can identify if two proteins interact and the accuracy of their structure. Never before has the potential for expanding the known structural understanding of protein interactions been this large, at such a small cost. There are currently 11.9 million pairwise human protein interactions in the String DB^32^. If 31% of these can be predicted at an error rate of 1%, this results in the structure of 3.7 million human heterodimeric protein structures. The computational cost to run all of this would take approximately three months on an Nvidia A100 system.

## Methods

### Development set

A set of heterodimeric complexes from Dockground benchmark 4^33^ is used to develop the pipeline, focusing on the AF2 configuration presented here. This set contains protein pairs, with each chain having at least 50 residues, sharing less than 30% sequence identity and no crystal packing artefacts. There are 219 protein interactions for which both unbound (singlechain) and bound (interacting chains) structures are available. Unbound chains share at least 97% sequence identity with the bound counterpart and, to facilitate comparisons, nonmatching residues are deleted and renumbered to become identical to the unbound counterpart. AF2 MSAs could not be generated for three of the complexes due to memory limitations (1gg2, 2nqd and 2xwb) using a computational node with 128 Gb RAM for the MSA generation and were thus disregarded, resulting in a total of 216 complexes. The dataset consists of 54% Eukaryotic proteins, 38% Bacterial and 8% from mixed kingdoms, e.g. one bacterial protein interacting with one eukaryotic.

### Test set

We used 1,661 protein complexes with known interfaces from_a recent study^27^ to test the developed pipeline. Here, three large biological assemblies were excluded. These complexes share less than 30% sequence identity, have a resolution between 1-5 Å and constitute unique pairs of PFAM domains (no single protein pair have PFAM domains matching that of any other pair). Some structures failed to be modelled for various reasons (see limitations of data generation), resulting in a total of 1481 structures. These proteins are mainly from H. Sapiens (25%), *S. Cerevisiae* (10%), *E.coli* (5%) and other Eukarya (30%).

107 of the complexes in the test set lack beta carbons (Cβs), and 50 have overlapping PDB codes with the development set and were therefore excluded. In the MSA generation from AF2, 20 MSAs report MergeMasterSlave errors regarding discrepancies in the number of match states, resulting in a total of 1484 AF2 MSAs. When folding, three of these (5AWF_D-5AWF_B, 2ZXE_B-2ZXE_A and 2ZXE_A-2ZXE_G) report “ValueError: Cannot create a tensor proto whose content is larger than 2GB”, leading to a final set of 1481 complexes. DSSP could only be run successfully for 1391 out of the 1481 protein complexes, and we ignored the rest in the analysis.

For RF, 26 complexes produced out of memory exceptions during prediction using a GPU with 40 Gb RAM and were excluded from the RF analyses, leaving 1455 complexes.

For the mammalian proteins from Negatome, seven out of 1733 single chains were redundant according to Uniprot (C4ZQ83, I0LJR4, I0LL25, K4CRX6, P62988, Q8NI70, Q8T3B2), 34 had no matching species in the MSA pairing, 106 produced out of memory exceptions during prediction using a GPU with 40 Gb RAM, 35 gave a tensor reshape error, and 65 complexes were homodimers, leaving 1715 complexes for this set.

### CASP14 set and novel protein complexes

As an additional test set, we used a set of six heterodimers from the CASP14 experiment. In addition, we extracted eight novel protein complexes deposited in PDB after 15 June 2021, which produced no results for at least one chain in each complex when submitted to the HHPRED web server (version 01-09-2021)^34,35^, see Table S1. We selected this small set to test the performance on data AF2 is guaranteed not to have seen.

### Non-interacting proteins

Two datasets of known non-interacting proteins were used, one from the same study as the positive test set^27^. Here, all proteins are from *E.coli.* Two methods were used to identify noninteracting proteins, first a set of proteins with no reported interaction signal in Yeast Two-Hybrid Experiments^36^ and secondly complexes whose individual proteins were found in different APMS benchmark complexes^37^. This dataset contains in total 3989 non-interacting pairs.

The second set contains 1964 unique mammalian protein complexes filtered against the IntAct^38^ dataset from Negatome^39^. This data deemed “the manual stringent set” contains proteins annotated from the literature with experimental support describing the lack of protein interaction. Some structures in this dataset are homodimers (65) and are therefore excluded, resulting in 1705 structures. Together there are 5694 non-interacting protein complexes.

### AlphaFold2 default MSA generation methodology

The input to AlphaFold2 (AF2) consists of several MSAs. We used the AF2 MSA generation^16^, which builds three different MSAs generated by searching the Big Fantastic Database^40^ (BFD) with HHBlits^41^ (from hh-suite v.3.0-beta.3 version 14/07/2017) and both MGnify v.2018_12^42^ and Uniref90 v.2020_01^43^ with jackhmmer from HMMER3^44^. The AF2 MSAs were generated by supplying a concatenated protein sequence of the entire complex to the AF2 MSA generating pipeline in FASTA format. The resulting MSAs will thus mainly contain gaps for one of the two query proteins in each row, as only single chains can obtain hits in the searched databases (Figure 6). No trimming or gap removal was performed on these MSAs.

**Figure 6.**
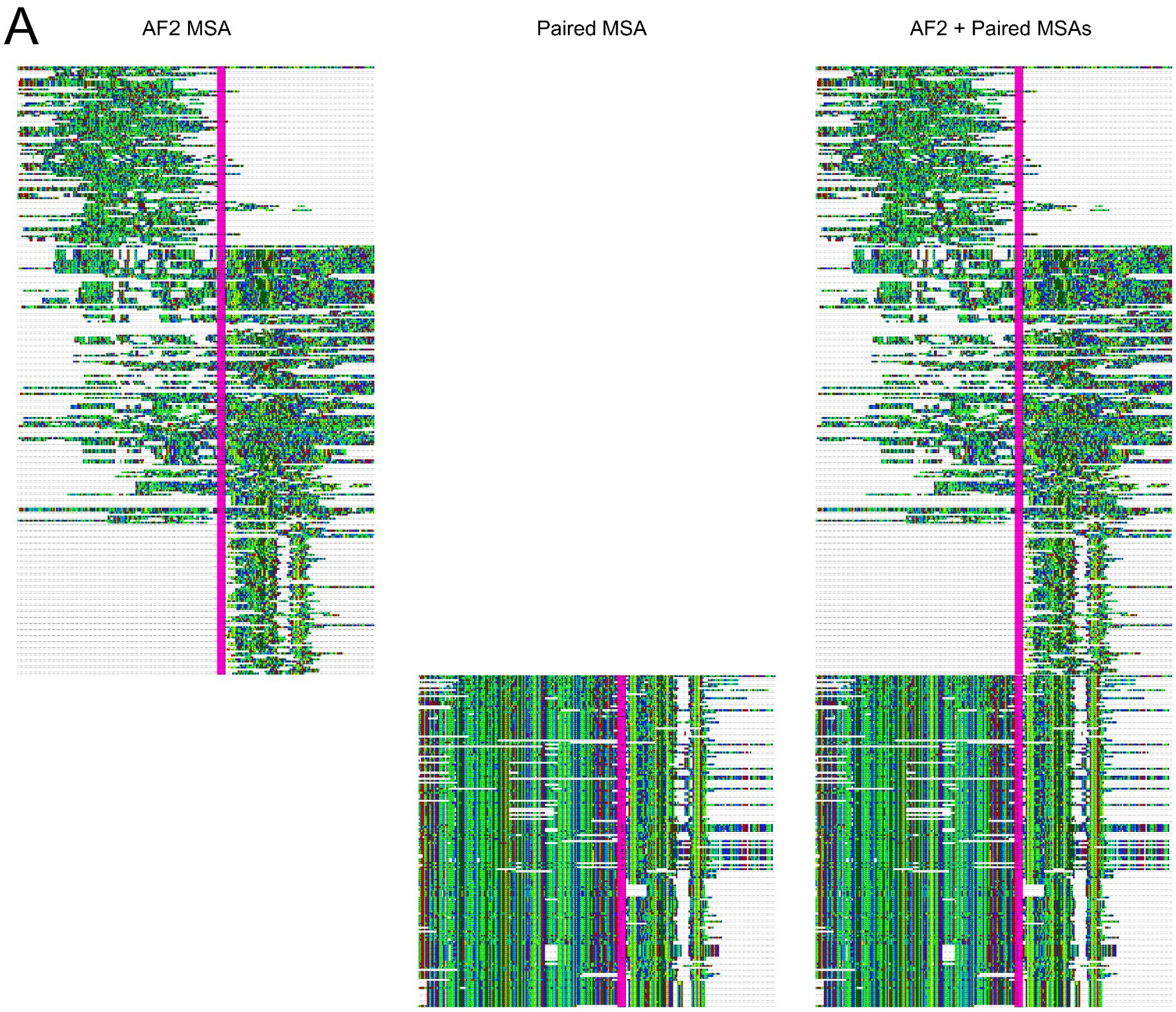
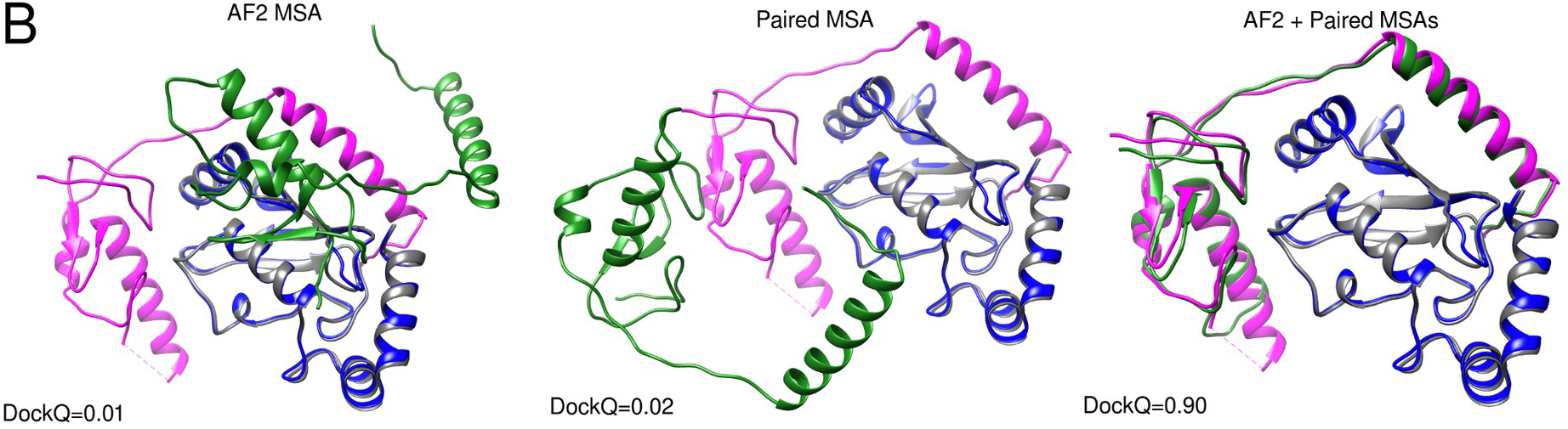
Comparison of different MSAs. **A)** Depiction of MSAs generated by AF2 and the paired version matched using organism information. Both AF and paired representations are sections containing 10% of the sequences aligned in the original MSA. Concatenated chains are separated by a vertical line (magenta). The visualisations were made using Jalview version 2.11.1.4^47^ **B)** Docking visualisations for PDB ID 5D1M with the model/native chains A in blue/grey and B in green/magenta using the three different MSAs in A. The DockQ scores are 0.01, 0.02 and 0.90 for AF2, paired, and AF2+paired MSAs, respectively.

### MSA block diagonalization

In addition to the default AF2 MSA, we generated an additional MSA by simply concatenating diagonally MSAs generated independently from each of the two chains. These MSAs were constructed by running HHblits^41^ version 3.1.0 against uniclust30_2018_08^45^ with these options:

~~~
hhblits -E 0.001 -all -oa3m -n 2
~~~

The concatenation is done by joining side-by-side the two input chains; then sequences from one MSA are added, aligned to the corresponding input chain. Each sequence in the MSA is then elongated with gaps (on the right side if it is the left sequence MSA or the other way around), to reach the length of the two concatenated input chains. The process is then repeated for the other input chain MSA to complete the block diagonalization.

### Paired MSA generation

In addition to the block diagonalization MSAs, we used a “paired MSA”, constructed using organism information, as described before^4,21,24^ (Figure 6). The rationale behind using a paired MSA is to identify inter-chain co-evolutionary information. An unpaired MSA has a limited inter-chain signal since the chains are treated in isolation.

The organism information was, using the OX identifier, was extracted from the two HHblits MSAs^46^. Next, all hits with more than 90% gaps were removed. From all remaining hits in the two MSAs, the highest-ranked hit from one organism was paired with the highest-ranked hit of the interacting chain from the same organism. Pairing the correct sequences should result in MSAs containing inter-chain co-evolutionary information^27^.

### Number of effective sequences (Neff)

To estimate the information in each MSA, we clustered sequences at 62% identity, as described in a previous study^48^. The number of clusters obtained in this way has been used to indicate a N_eff_ value for each MSA.

Unaligned FASTA sequences were extracted from the three AF2 default MSAs. Obtained sequences were processed with the CD-HIT software^49^ version 4.7 (http://weizhong-lab.ucsd.edu/cd-hit/) using the options:

~~~
-c 0.62 -G 0 -n 3 -aS 0.9
~~~

We calculated the Neff scores separately for paired and AF2 MSAs.

### AlphaFold2

We modelled complexes using AlphaFold2^16^ (AF2) by modifying the script https://github.com/deepmind/alphafold/blob/main/run_alphafold.py to insert a chain break of 200 residues - as suggested in the development of RoseTTAFold^17^ (RF). During modelling, relaxation was turned off, and only the atoms generated in RF (N, CA, C) were used in subsequent analyses. Sidechains were thus not used to score interfaces. We note that performing model relaxation did not increase performance in the AF2 paper^16^ and was, therefore, ignored to save computational cost. No templates were used to build structures, as this would not assess the prediction accuracy of unknown structures or structures without sufficient matching templates. Further, AF2 has been shown to perform well for single chains without templates and has reported higher accuracy than template-based methods even when robust templates are available^16^.

We supplied three different types of MSAs to AF2: the MSAs generated by using the default AF2 settings, the top paired MSAs constructed using HHblits, described above, and finally, a concatenation of these both alignments. AF2 was run with two different network models, AF2 model_1 (used in CASP14) and AF2 model_1_ptm, for each MSA. The second model, model_1_ptm, is a fine-tuned version of model_1 that predicts the TMscore^50^ and alignment errors^16^. We ran these two different models by using two different configurations. The configurations utilise a varying amount of recycles and ensemble structures. Recycle refers to the number of times iterative refinement is applied by feeding the intermediate outputs recursively back into the same neural network modules. At each recycling, the MSAs are resampled, allowing for new information to be passed through the network. The number of ensembles refers to how many times information is passed through the neural network before it is averaged^16^. The two configurations used are; the CASP14 configuration (three recycles, eight ensembles) and an increased number of recycles (ten) but only one ensemble.

Since structure prediction with AF2 is a non-deterministic process, we generate five models initiated with different seeds. To save computational cost, this was only performed for the best modelling strategy. We rank the five models for each complex by the number of residues in the interface, giving the best result.

### RoseTTAFold

For comparison, the RoseTTAFold (RF) end-to-end version^17^ was run using the paired MSAs with the top hits. The RoseTTAFold pipeline for complex modelling only generates MSAs for bacterial protein complexes, while the proteins in our test set are mainly Eukaryotic. Therefore, we use the paired alignments here. We compare RF with AF2 using the same inputs (the paired MSAs) for both the development and test datasets to provide a more fair comparison, as AF2 searches many different databases to obtain as much evolutionary information as possible when generating its MSAs. To predict the complexes, we use the “chain break modelling” as suggested in RF (https://github.com/RosettaCommons/RoseTTAFold/tree/main/example/complex_modeling) using the following command:

~~~
predict_complex.py -i msa.a3m -o complex -Ls chain1_length chain2_length
~~~

No optimisation of the RF protocol was made here.

### MDockPP

The docking method MDockPP^30^ was run through the provided webserver (https://zougrouptoolkit.missouri.edu/MDockPP/). This docking algorithm is based on Fast Fourier Transform (FFT). The docking results are assessed using the “in-house” scoring function ITScorePP.

### GRAMM

For comparison, a rigid-body docking method, GRAMM^31^, was used. Here, two protein models are docked using a Fast Fourier Transform (FFT) procedure to generate 340’000 docking poses for each complex. The bound structures extracted from complexes in the test set were used as inputs. This docking generation stage mainly considers the geometric surface properties of the two interacting structures, allowing minor clashes to leave some space for conformational flexibility adjustment. As the bound form of the proteins is used, this should represent an easy case for GRAMM based docking, and performance drops significantly when unbound structures or models are used ^51^. The atom-atom contact energy AACE18 is used to score and rank all poses, as this has been shown to provide better results than shape-complementarity alone^52^.

### Template-based docking

For comparison, a template-based docking protocol^7^ referred to as “TMdock” is also adopted. The adopted template library includes 11756 protein complexes obtained from the Dockground database^33^ (release 28-10-2020). Monomers from target complexes are structurally aligned with complexes in the supplied libraries (depleted of the target structure PDB ID) in order to identify the best available template structure. The bound form of the template structures was used. TM-scores resulting from the alignment of target proteins to each template are averaged and used to score obtained docking models. Alternatively, we refer to “TMdock Interfaces” when targets are structurally aligned only to the template interfaces, defined as every residue with a Cβ atom closer than 12 Å from any Cβ atom in the other chain.

### AlphaFold-multimer

The simultaneous fold-and-dock program based on the same principles as AF2, AlphaFold-multimer^28^, was run with the default settings. These entail creating four different MSAs. Three different MSAs are created by searching Uniref90 v.2020_01^43^, Uniprot v.2021_04^46^ and MGnify v.2018_12^42^ with jackhmmer from HMMER3^44^ and one joint is created by searching the Big Fantastic Database^40^ (BFD) and uniclust30_2018_08^45^ with HHBlits^41^ (from hh-suite v.3.0-beta.3 version 14/07/2017).

The results from the Uniprot search are used for MSA pairing and the results from the remaining searches are used to create a block-diagonalized MSA, similar to the procedures described above. All four MSAs are then used to fold a protein complex. Some complexes failed due to computational limitations, resulting in 1458 out of 1481 complexes successfully folded.

### Scoring models

The backbone atoms (N, CA and C) were extracted from the predicted AF2 structures (as these are the only predicted atoms in the end-to-end version of RF). The interface scoring program DockQ^53^ was then run (without any special settings) to compare the predicted and actual interfaces. This program compares interfaces using a combination of three different CAPRI^54^ quality measures (F_nat_, LRMS, and iRMS) converted to a continuous scale, where an acceptable model comprises a DockQ score of at least 0.23.

### Ranking models

To analyse the ability of AF2 to distinguish correct models as well as interacting from non-interacting proteins, we analyse the separation between acceptable and incorrect models as a function of different metrics on the development set: the number of unique interacting residues (Cβs from different chains within 8 Å from each other), the total number of interactions between Cβs from different chains (referred to as the number of interface contacts), average predicted lDDT (plDDT) score from AF2 for the interface, the minimum of the average plDDT for both chains and the average plDDT over the whole heterodimer.

We use these metrics as a threshold to build a confusion matrix, where True/False Positives (TP and FP respectively) are correct/incorrect docking models which places above the threshold and False/True Negatives (FN and TN respectively) are correct/incorrect docking models which scores below the threshold. From the built confusion matrix, we derive the True Positive Rate (TPR), False Positive Rate (FPR) defined as:

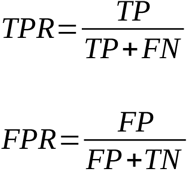

Then, we calculate TPR and FPR for each possible value assumed by the set of dockings given a single metric and plot TPR as a function of FPR in order to obtain a Receiver Operating Characteristic (ROC) curve. We compute the Area Under Curve (AUC) for ROC curves obtained for each metric to compare different metrics. The AUC is defined as:

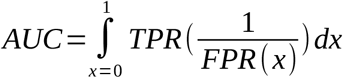

The TPR and FPR for different thresholds are used to calculate the fraction of models that can be called correct out of all models and the Positive Predictive Value (PPV). The fraction of acceptable and incorrect models are obtained by multiplying the TPR and FPR with the success rate (SR). Multiplying the FPR with the SR results in the False Discovery rate(FDR) and the PPV can be calculated by dividing the fraction of acceptable models by the sum of the acceptable and incorrect models. The PPV, FDR and SR are defined as:

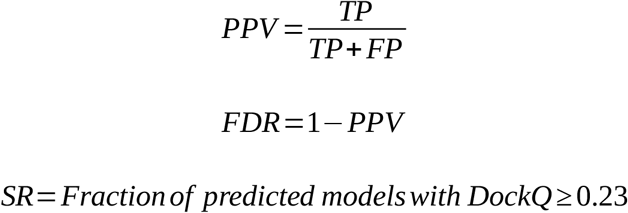

### pDockQ

As it is not only desirable to know when a model is accurate but also how accurate this model is, we developed a predicted DockQ score, pDockQ. This score is created by fitting a sigmoidal curve (Figure 2C) using “curve_fit” from SciPy v.1.4.1 ^55^, to the DockQ scores using the IF plDDT·log(IF contacts), with the following sigmoidal equation: where x = average interface plDDT·log(number of interface contacts) and we obtain L= 0.958, X_0_ = 160.11, k = 1 and b = 0.001.

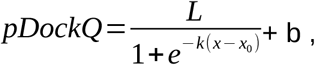

### Analysis of models

To analyse the possibility of determining when AF2 can model a complex correctly, we analyse the structures and the multiple sequence alignments. We investigated: the number of effective sequences (Neff), the secondary structure in the interface annotated using DSSP^56^, the length of the shortest chain, the number of residues in the interface and the number of contacts in the interface.

DSSP was run on the entire complexes, and the resulting annotations were grouped into three categories; helix (3-turn helix (3_10_ helix), 4-turn helix (α helix) and 5-turn helix (π helix)), sheet (extended strand in parallel or antiparallel β-sheet conformation and residues in isolated β-bridges) and loop (residues which are not in any known conformation).

In addition, we assess the positive predictive value (PPV) of the top N interface direct coupling analysis (DCA) signals using the paired MSAs. Here, N is the number of true interface contacts (Cβs from different chains within 8 Å from each other). The PPV is therefore the fraction of the top N DCA signals in the interface that are true contacts. The DCA signals are computed using GaussDCA^57^.

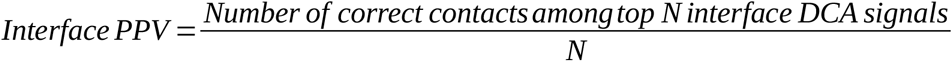

### Computational cost

To compare the computation required for each MSA, we compared the time it took to generate MSAs for three protein pairs (PDB: 4G4S_P-O, 5XJL_A-2 and 5XJL_2-M), using either the block diagonalization or AF2 protocol. The tests were performed on a computer using 16 CPU cores from an Intel Xeon E5-2690v4.

Fusing the MSAs took 3 seconds on average per tested complex. It took 7884 seconds for generating the AF2 MSAs, the single-chain searches took 338 seconds on average and the pairing 2 seconds. The pairing and fusing are thereby negligible compared to searching, resulting in a speedup of 24 times for the hhblits searches. In comparison, folding using the m1-10-1 strategy took 191 seconds on average for these pairs.

## Data availability

All data used to produce the results are available at: https://doi.org/10.17044/scilifelab.16866202.v1;

Results used to produce all figures can be found in the supplementary information.

## Code availability

All code to run FoldDock and reproduce the analysis here can be obtained here https://gitlab.com/ElofssonLab/FoldDock (commit 2e4c96aa352338976260ece0646ceaaa75392dec) under the Apache License, Version 2.0.

## Acknowledgements

We thank Petras Kundrotas for supplying the new heterodimeric proteins without templates in the PDB. We also thank Liming Qiu and Xiaoqin Zou for their help with running their docking program MDockPP in a timely manner.

Financial support: Swedish Research Council for Natural Science, grant No. VR-2016-06301 and Swedish E-science Research Center. Computational resources: Swedish National Infrastructure for Computing, grants: SNIC 2021/5-297, SNIC 2021/6-197 and Berzelius-2021-29.

## Author contributions

PB and GP performed the studies; all authors contributed to the analysis. PB wrote the first draft of the manuscript; all authors contributed to the final version. AE obtained funding.

## Supplementary Material

### Tables

**Table S1.** Results over CASP14 and new heterodimers

| Table S1. Results over CASP14 and new heterodimers |  |  |  |
| --- | --- | --- | --- |
| PDB | Chain 1 | Chain 2 | Top-ranked DockQ score |
| CASP 14 heterodimers |  |  |  |
| 7M5F | A | C | 0.85 |
| 6XOD | A | B | 0.94 |
| 6VN1 | A | H | 0.01 |
| 6VN1 | H | L | 0.45 |
| 6R17 | A | C | 0.54 |
| 6TMM* | A | B | 0.03 |
| 6TMM* | A | D | 0.04 |
| Heterodimers without templates |  |  |  |
| 7EIV | A | C | 0.76 |
| 7EL1 | A | E | 0.01 |
| 7K01 | 1 | 6 | 0.36 |
| 7LDG | A | B | 0.45 |
| 7LF7 | A | M | 0.02 |
| 7LF7 | B | M | 0.02 |
| 7MEZ | A | B | 0.53 |
| 7NJ0 | A | C | 0.04 |

The two interactions involving 6TMM are different possible configurations of the same two chains.

**Table S2.**
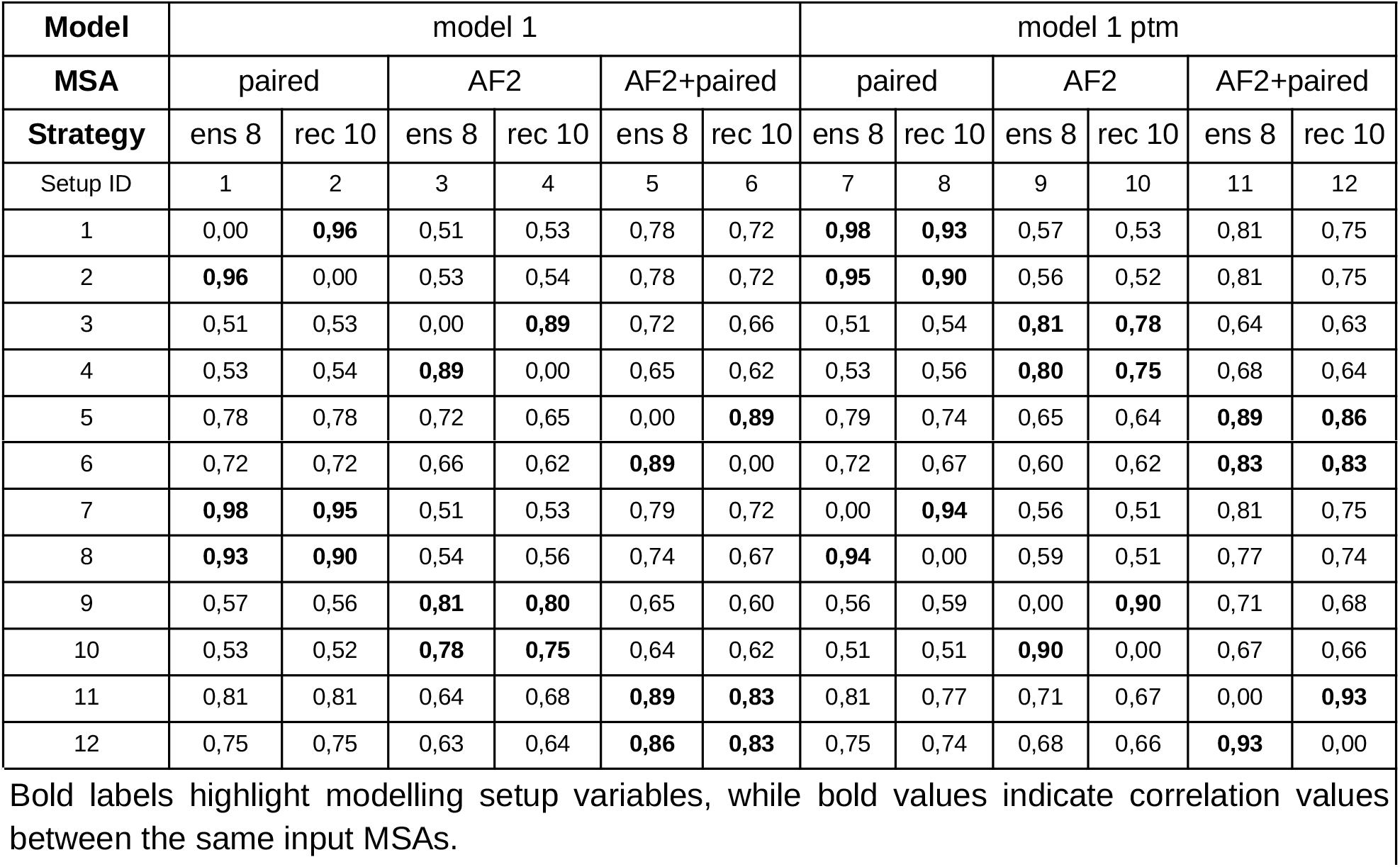
Pearson correlation coefficients of DockQ sets from different modelling setup

The two interactions involving 6TMM are different possible configurations of the same two chains.
|  |  |  |  |  |  |  |  |  |  |  |  |  |
| --- | --- | --- | --- | --- | --- | --- | --- | --- | --- | --- | --- | --- |
| Model | model 1 |  |  |  |  |  | model 1 ptm |  |  |  |  |  |
| MSA | paired |  | AF2 |  | AF2+paired |  | paired |  | AF2 |  | AF2+paired |  |
| Strategy | ens 8 | rec 10 | ens 8 | rec 10 | ens 8 | rec 10 | ens 8 | rec 10 | ens 8 | rec 10 | ens 8 | rec 10 |
| Setup ID | 1 | 2 | 3 | 4 | 5 | 6 | 7 | 8 | 9 | 10 | 11 | 12 |
| 1 | 0,00 | **0,96** | 0,51 | 0,53 | 0,78 | 0,72 | **0,98** | **0,93** | 0,57 | 0,53 | 0,81 | 0,75 |
| 2 | **0,96** | 0,00 | 0,53 | 0,54 | 0,78 | 0,72 | **0,95** | **0,90** | 0,56 | 0,52 | 0,81 | 0,75 |
| 3 | 0,51 | 0,53 | 0,00 | **0,89** | 0,72 | 0,66 | 0,51 | 0,54 | **0,81** | **0,78** | 0,64 | 0,63 |
| 4 | 0,53 | 0,54 | **0,89** | 0,00 | 0,65 | 0,62 | 0,53 | 0,56 | **0,80** | **0,75** | 0,68 | 0,64 |
| 5 | 0,78 | 0,78 | 0,72 | 0,65 | 0,00 | **0,89** | 0,79 | 0,74 | 0,65 | 0,64 | **0,89** | **0,86** |
| 6 | 0,72 | 0,72 | 0,66 | 0,62 | **0,89** | 0,00 | 0,72 | 0,67 | 0,60 | 0,62 | **0,83** | **0,83** |
| 7 | **0,98** | **0,95** | 0,51 | 0,53 | 0,79 | 0,72 | 0,00 | **0,94** | 0,56 | 0,51 | 0,81 | 0,75 |
| 8 | **0,93** | **0,90** | 0,54 | 0,56 | 0,74 | 0,67 | **0,94** | 0,00 | 0,59 | 0,51 | 0,77 | 0,74 |
| 9 | 0,57 | 0,56 | **0,81** | **0,80** | 0,65 | 0,60 | 0,56 | 0,59 | 0,00 | **0,90** | 0,71 | 0,68 |
| 10 | 0,53 | 0,52 | **0,78** | **0,75** | 0,64 | 0,62 | 0,51 | 0,51 | **0,90** | 0,00 | 0,67 | 0,66 |
| 11 | 0,81 | 0,81 | 0,64 | 0,68 | **0,89** | **0,83** | 0,81 | 0,77 | 0,71 | 0,67 | 0,00 | **0,93** |
| 12 | 0,75 | 0,75 | 0,63 | 0,64 | **0,86** | **0,83** | 0,75 | 0,74 | 0,68 | 0,66 | **0,93** | 0,00 |

Bold labels highlight modelling setup variables, while bold values indicate correlation values between the same input MSAs.

**Table S3.** Quality metrics for test set AF2 ranked models

| Table S3. Quality metrics for test set AF2 ranked models |  |  |  |  |  |
| --- | --- | --- | --- | --- | --- |
| TPR | FPR | PPV | Fraction Correct | Fraction Incorrect | pDockQ |
| 0.370 | 0.007 | 0.981 | 0.228 | 0.004 | 0.674 |
| 0.416 | 0.016 | 0.963 | 0.257 | 0.010 | 0.655 |
| 0.504 | 0.025 | 0.953 | 0.311 | 0.015 | 0.629 |
| 0.525 | 0.034 | 0.940 | 0.324 | 0.021 | 0.622 |
| 0.578 | 0.042 | 0.932 | 0.357 | 0.026 | 0.596 |
| 0.624 | 0.051 | 0.924 | 0.385 | 0.032 | 0.577 |
| 0.656 | 0.060 | 0.916 | 0.405 | 0.037 | 0.558 |
| 0.691 | 0.069 | 0.910 | 0.427 | 0.042 | 0.535 |
| 0.718 | 0.079 | 0.900 | 0.443 | 0.049 | 0.516 |
| 0.757 | 0.092 | 0.892 | 0.467 | 0.057 | 0.490 |
| 0.779 | 0.101 | 0.886 | 0.481 | 0.062 | 0.471 |
| 0.814 | 0.113 | 0.878 | 0.502 | 0.070 | 0.441 |
| 0.835 | 0.123 | 0.871 | 0.515 | 0.076 | 0.421 |
| 0.848 | 0.138 | 0.860 | 0.523 | 0.085 | 0.396 |
| 0.860 | 0.146 | 0.855 | 0.531 | 0.090 | 0.381 |
| 0.871 | 0.162 | 0.843 | 0.537 | 0.100 | 0.362 |
| 0.884 | 0.176 | 0.834 | 0.546 | 0.109 | 0.344 |
| 0.898 | 0.194 | 0.822 | 0.554 | 0.120 | 0.326 |
| 0.906 | 0.210 | 0.812 | 0.559 | 0.130 | 0.315 |
| 0.916 | 0.226 | 0.802 | 0.565 | 0.139 | 0.298 |
| 0.930 | 0.254 | 0.785 | 0.574 | 0.157 | 0.269 |
| 0.938 | 0.268 | 0.778 | 0.579 | 0.165 | 0.249 |
| 0.946 | 0.300 | 0.759 | 0.584 | 0.185 | 0.223 |
| 0.954 | 0.335 | 0.740 | 0.589 | 0.207 | 0.201 |
| 0.961 | 0.354 | 0.730 | 0.593 | 0.219 | 0.181 |
| 0.968 | 0.388 | 0.714 | 0.598 | 0.239 | 0.160 |
| 0.975 | 0.406 | 0.706 | 0.602 | 0.250 | 0.145 |
| 0.980 | 0.448 | 0.686 | 0.605 | 0.276 | 0.126 |
| 0.987 | 0.492 | 0.667 | 0.609 | 0.304 | 0.106 |
| 0.993 | 0.570 | 0.636 | 0.613 | 0.352 | 0.074 |
| 0.999 | 0.788 | 0.559 | 0.616 | 0.487 | 0.030 |

### Figures

**Figure S1.**
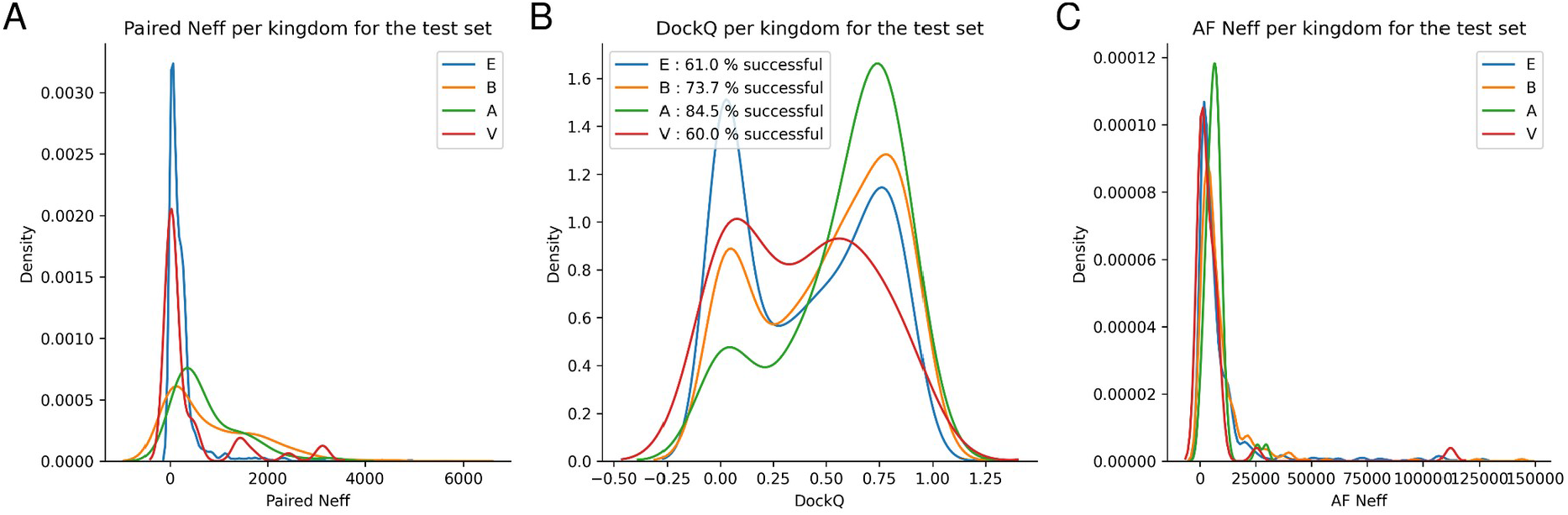
Distribution of different test set features divided by kingdom. In the figure different features **A)** Paired Neff, **B)** DockQ scores, and **C)** AF Neff are plotted according to complex provenience from: E=Eukarya, B=Bacteria, A=Archaea and V=Virus.

**Figure S2.**
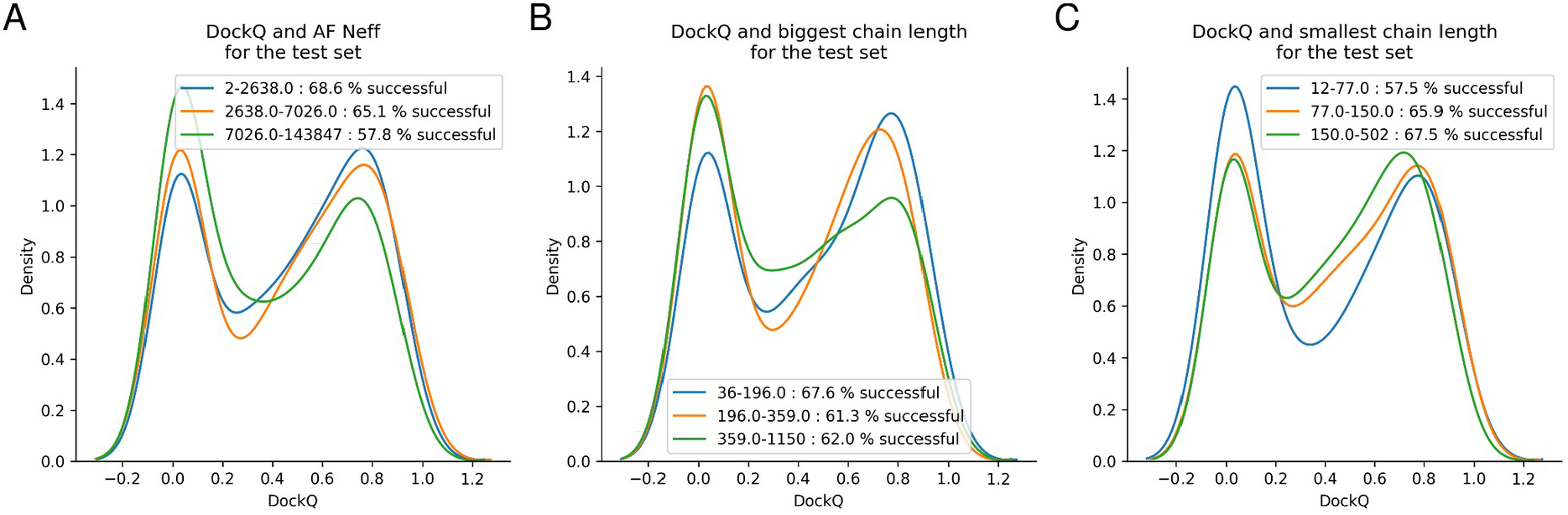
DockQ distributions for test dataset tertiles. Tertiles are derived from **A)** AF Neff **B)** biggest chain lengths and **C)** smallest chain length. The separation between the tertiles is low for all features displaying similar success rates.

**Figure S3.**
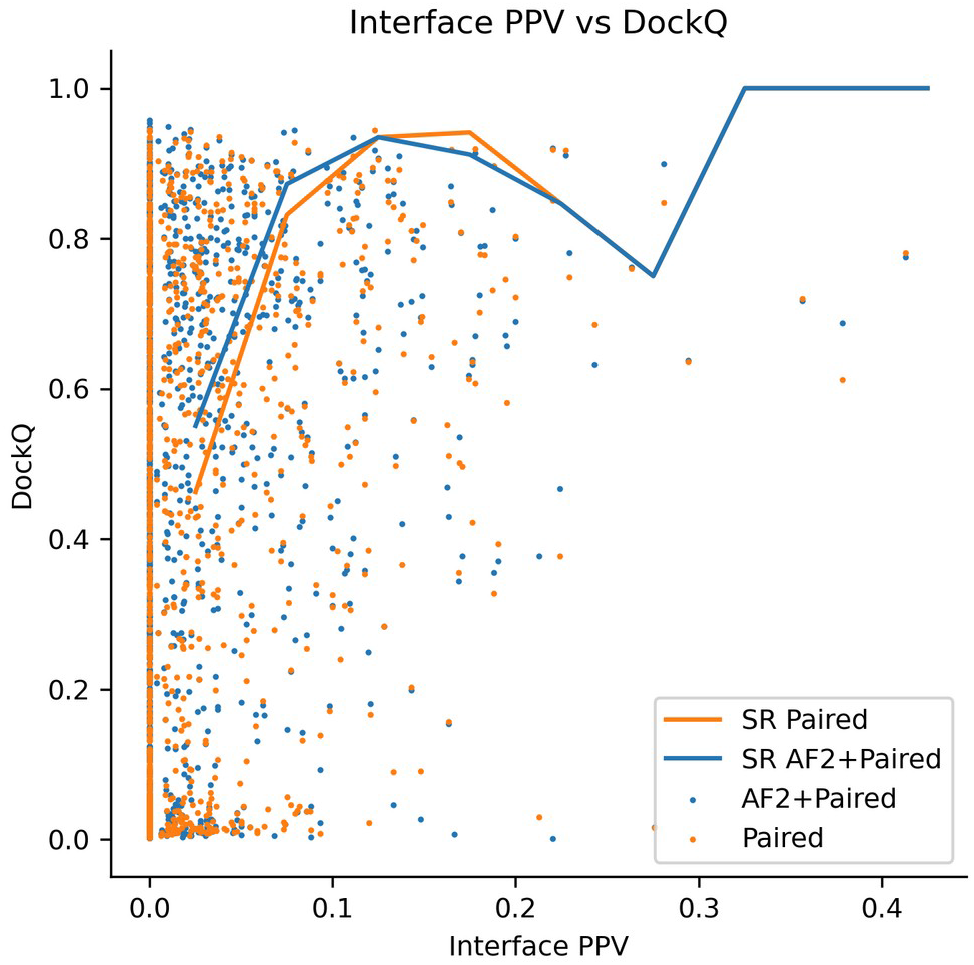
Interface PPV in paired MSAs vs DockQ scores and success rates (SR). The SR increases with interface PPV for both paired and AF2+paired modelings, suggesting a strong influence of the MSAs on the outcome.

**Figure S4.**
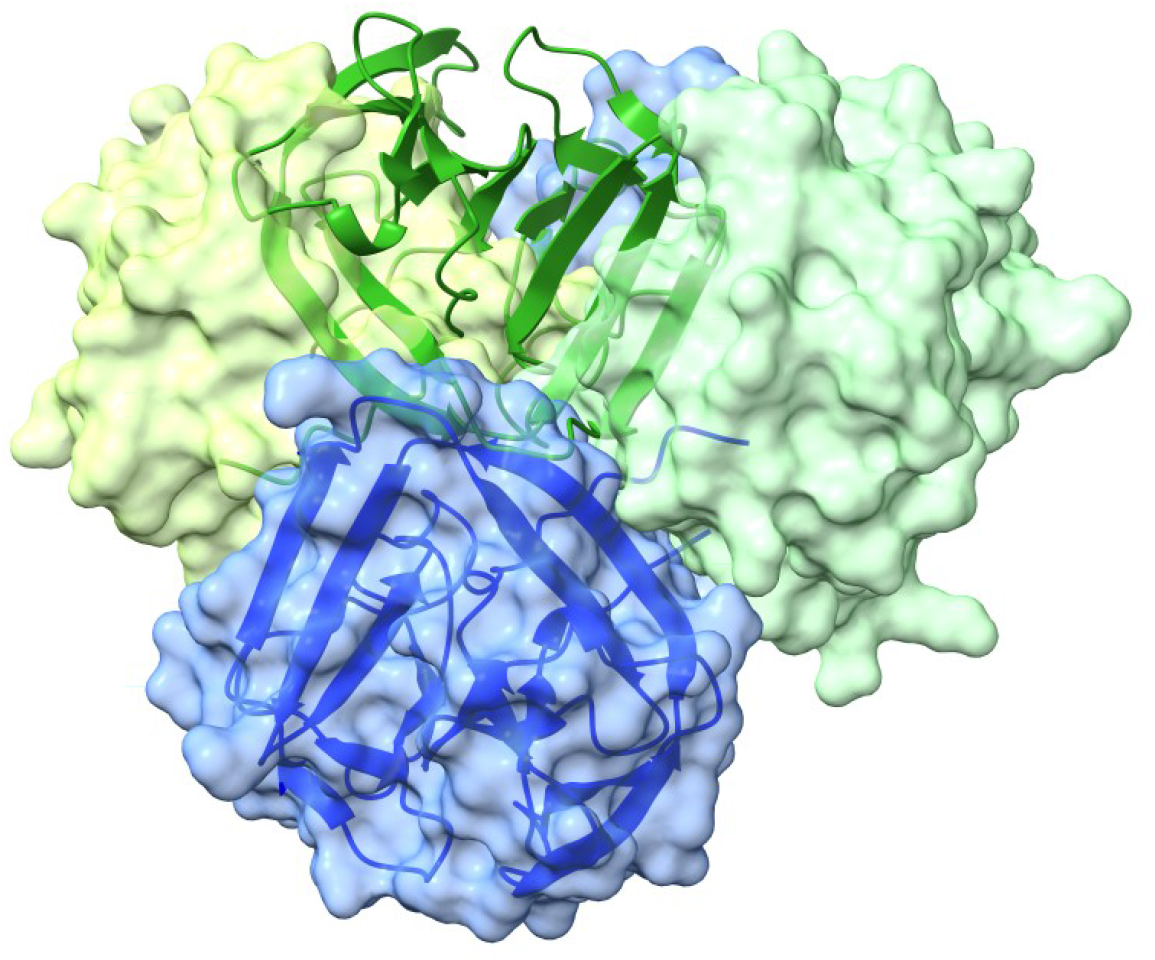
Model of CASP14 heterodimer from PDB 6TMM. The model obtained using AF2 default and paired MSA (ribbons) is superposed to the native heterocomplex (surfaces). The docking model smaller chain (green ribbon) is positioned halfway between the two alternative binding sites formed by blue and green surfaces.

**Figure S5.**
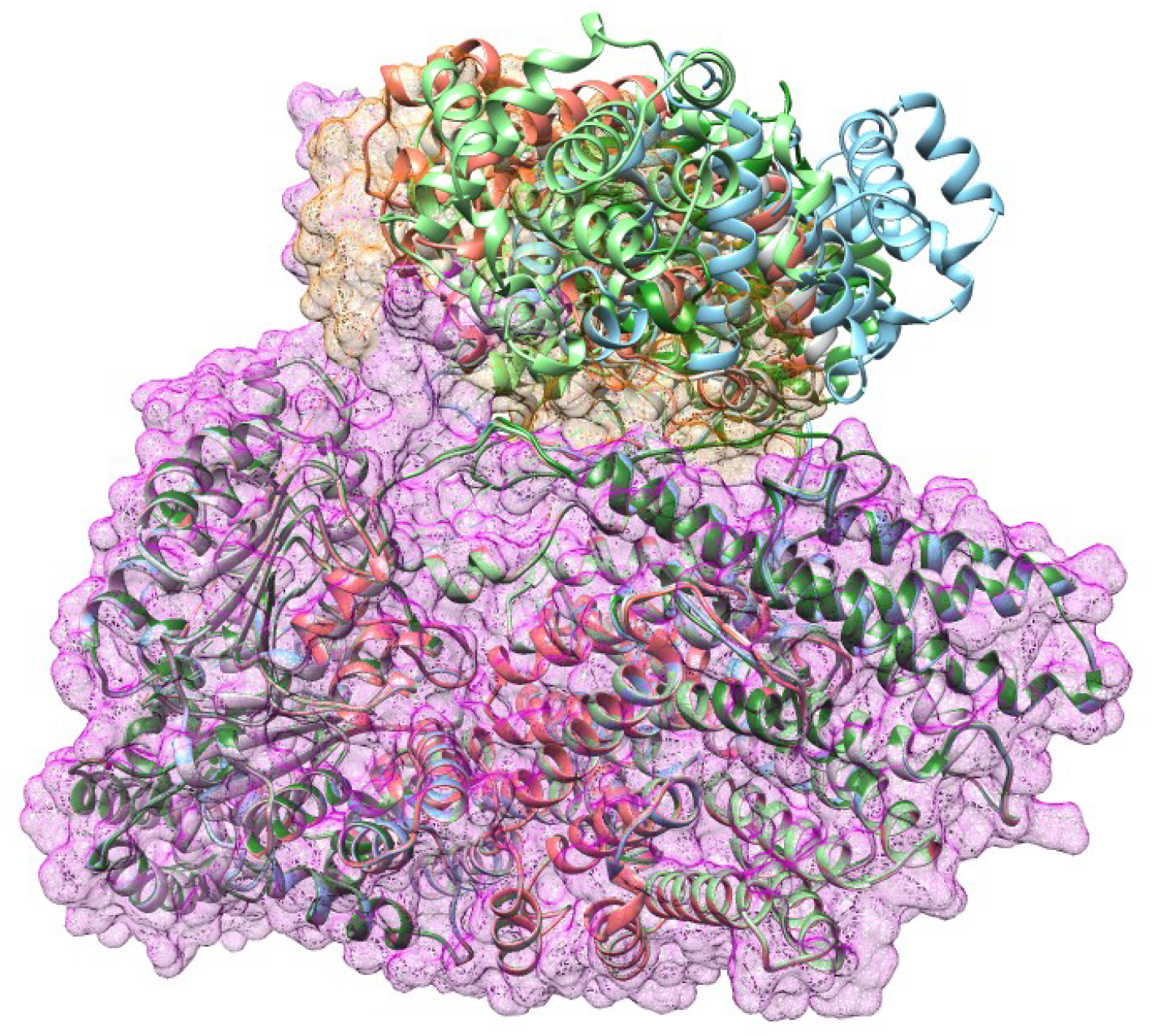
Model of heterodimer from PDB 7NJ0. The native structure is represented as a mesh surface (orange and magenta). All predictions (ribbons) get the location of the chains correct, but the interface and orientations are slightly wrong, resulting in DockQ scores close to 0.

